# Wearable Assessment of Level and Uphill Running at Critical Intensity

**DOI:** 10.1101/2025.04.25.650654

**Authors:** Nikolai Kirner, Wouter Hoogkamer

**Affiliations:** Integrative Locomotion Laboratory, Department of Kinesiology, University of Massachusetts, Amherst, MA, USA; Department Health and Sport Sciences, TUM School of Medicine and Health, Technical University of Munich, Munich, Germany

**Keywords:** Critical Power, Critical Speed, Time-to-exhaustion, Running Power, 3-minute all-out test, Trail Running

## Abstract

**Purpose:** We used commercially available wearable devices that measure running power (foot- vs. wrist-based) and muscle oxygenation to investigate time-to-exhaustion during level and uphill running at critical intensity.

**Methods:** 14 healthy recreational runners participated in this study. They first ran a 3-minute all-out test to derive their critical running intensity. They then performed two time-to-exhaustion trials (TTEs) at this intensity at separate slopes to assess time-to-exhaustion, muscle oxygenation, and rating of perceived exertion.

**Results:** We observed a shorter time-to-exhaustion (539 ± 160 vs. 936 ± 336 s), a faster decline in muscle oxygenation over time (-2.48 ± 1.32 vs. -1.62 ± 2.11 %·min^-1^), a higher rating of perceived exertion at matched time points, and a higher estimated metabolic rate (19.51 ± 1.99 vs. 17.82 ± 1.62 W·kg^-1^) for the level time-to-exhaustion test (TTE_Level_) compared to the uphill time-to-exhaustion test (TTE_Incline_) at a 2.86° (5%) slope.

**Conclusion:** Our data suggests that athletes should be careful when using running power at critical intensity to guide their racing and training in non-level conditions.

## Introduction

In 1986, Ulrich Schoberer developed the first spider-based power meter for cycling to measure an athlete’s work rate during training and racing. The system revolutionized the sport by enabling cyclists to quantify their effort in a single metric independent of weather conditions, terrain, and physiological considerations using their own bikes. The spider-based power meter uses strain gauges to measure the force applied on the chainring during pedaling, considering the crank’s angular velocity (cadence) (Montalvo-Pérez et al. 2021). Further systems, such as pedal- or crank-based power meters, have expanded the technology over the last decades (Passfield et al. 2017) since early adopters such as the East German Cycling National Team and Tour de France Champions Greg LeMond started to use the system in the 1990s, with LeMond attributing his victories to the use of a power meter (SRM GmbH 2024).

Based on this success, engineers, inventors, and scientists have tried to develop a similar device for running that provides an effort metric going beyond heart rate or speed measurements, which are substantially affected by factors such as temperature (Rojas-Valverde et al. 2020), slope (Margaria et al. 1963; Brill and Kram 2021), physiological response times or caffeine intake (Turley 2016; Stadheim et al. 2021) to highlight a few. With the introduction of running power devices in the mid-2010s, estimating power is now accessible using commercially available sensors worn at the wrist (e.g., Garmin and Coros watches) or foot (e.g., Stryd and RunScribe). However, running power is more controversial than cycling power (van Rassel et al. 2023).

In cycling, riders produce mechanical power by applying torque on the crank, achieving rotational velocity (Passfield et al. 2017), which is highly correlated with metabolic rate (Gaesser and Brooks 1975). Any fluctuations in external power demands due to drivetrain efficiency, tire pressure, road surface, air resistance, or slope at a constant power output will affect cycling speed, but not metabolic rate. Hence, a rider’s power output reflects their metabolic effort to reach desired speeds. Contrarily, in running, mechanical power output is poorly correlated to metabolic rate (Heglund et al. 1982). Running energetics are characterized by negative work during the first half of the stance phase, followed by positive work during the second half of the stance phase (Cavagna et al. 1964), but most of that work is performed through elastic strain energy storage and return in passive tissues rather than by active muscle fiber work (Fenn 1930; Cavagna and Kaneko, 1977; Lai et al. 2014; Swinnen et al. 2019; Monte et al. 2020). To keep these elastic tissues under tension, muscles must produce force isometrically without performing work (Fenn 1930). Therefore, the metabolic cost of running is dominated by the metabolic cost of generating force (Kram and Taylor, 1990). Furthermore, while the positive work in cycling is done by active muscle fiber work at a fairly constant efficiency (e.g., ranging from ∼20-22% in elite cyclists (Jeukendrup et al. 2003)), the running efficiency of any actively performed mechanical work will differ between positive and negative work (Margaria 1968). These differences in elastic energy storage and return and varying efficiencies for positive vs. negative work complicate the use of mechanical power to assess metabolic effort in running.

To circumvent these challenges, commercially available running sensors use proprietary algorithms to estimate the positive mechanical work and the associated metabolic energy expenditure. One of the most popular devices is Stryd’s foot pod (Stryd Inc., Boulder, CO, USA), which uses a 6-axis inertial motion sensor (3-axis gyroscope and 3-axis accelerometer). Stryd’s internal testing suggests that their mechanical power output estimation (W·kg^-1^) is highly correlated to oxygen uptake (ml·min^-1^·kg^-1^) (R^2^ = 0.96), allegedly confirmed by third-party research (Stryd Team 2017). Peer-reviewed research has reported more mixed and controversial results (see Aubry et al. (2018) and Snyder et al. (2018)). Taboga et al. (2022) reported that the sensor’s power estimates correlate with force plate-based power and metabolic rate across speeds for level treadmill running. Similarly, Imbach et al. (2020) observed correlations between Stryd running power and oxygen uptake for six individuals over a range of speeds during overground running. For running power to be more beneficial than running speed for measuring running intensity, validity must be assessed beyond speed dependence. Unfortunately, Stryd’s running power appears to be not adequately sensitive to differences in metabolic effort between advanced footwear technology and traditional running shoes (Hudgins et al. 2024). Furthermore, the validity of running power during uphill and downhill running is widely overlooked in the current literature (Apte et al. 2023). Altogether, it remains to be determined whether the Stryd Food Pod can accurately reflect changes in metabolic rate due to differences in running conditions other than speed (Austin et al. 2018).

Interestingly, the Stryd foot pod has been extensively tested to estimate critical power (CP) and critical speed (CS) using time-trial efforts (Dearing & Paton 2023; Hunter et al. 2023; van Rassel et al. 2024) or a 3-minute all-out test (3MT) (Hunter et al. 2023). CP estimations using Stryd’s foot pod have also been compared and validated to anaerobic threshold concepts (Ruiz-Alias et al. 2022; van Rassel et al. 2023). Hence, from a practical perspective, the validity of running power concerning metabolic rate in varying conditions appears secondary to its validity in providing consistent critical power estimates to inform training intensities and race strategies.

For this study, we aimed to investigated the usability of the Stryd foot pod to estimate running power for level and uphill running at critical intensity through assessing time-to-exhaustion, muscle oxygenation using near-infrared spectroscopy, rating of perceived exertion, and estimating metabolic rate.

## Methods

Fourteen healthy runners participated in the study (2 females; identified via self-reported sex assigned at birth; 70.7 ± 15.3 kg; 1.79 ± 0.13 m; 25.1 ± 5.7 years old). Participants gave their written informed consent before completing the first of three study visits as approved by the University of Massachusetts, Amherst Institutional Review Board (Protocol 5639). All study sessions, comprised of one session on the track and two lab sessions on the treadmill, were completed within 2 weeks, with a maximum of 1 week between each visit. Participants were recreational or trained runners who ran at least 10 miles weekly (up to 70 miles) with no lower limb injuries three months prior and during study participation.

### Session 1

Participants completed a physical activity readiness questionnaire (PAR-Q, Chisholm et al. 1975) before we recorded their body mass (portable scale) and height (self-reported). We took exact height measurements later at the start of the first lab visit (session 2). We placed a latest-generation Stryd foot pod centrally on the participant’s left shoelaces, according to the manufacturer’s guidelines. Additionally, participants wore a GPS running watch (Apex 2, Coros Wearables Inc. Shenzhen, China) on their left arm to record speed and watch-based running power with the display on the opposite side of their wrist. We set the participant’s mass and height before each trial using the Stryd App (Version 8.17.64) in Outdoor Run Mode and the Coros Watch in Track Mode. After a short device initialization, participants ran a self-paced 8-minute warmup, including two sub-maximal sprints of 20 meters. Following a passive rest of at least 5 minutes, participants performed a self-paced 3MT in lane 1. An examiner gave strong verbal encouragement to the participants riding alongside on a bike and ended their test after ∼3:10 minutes; a minimum test duration of 3:05 minutes ensured sufficient data sampling (Pettitt 2016). We calculated CS and CP as the average running speed and running power from the Stryd foot pod over the last 30 seconds of the 3MT (Hunter et al. 2023), starting 150 seconds after the first recorded data point for power, which matched the first recorded positive speed value. We also calculated and compared CS and CP for Coros’ Speed and Power data by calculating the average running speed and power of the last 30 seconds of each data set, starting with the first positive index of each speed and power vector (CS_Coros_ and CP_Coros_, respectively). CS marked the reference point for sessions 2 and 3. Before each 3MT, an examiner narrated an oral script with an analogy on how to approach the 3MT, enhancing compliance and standardizing the trial across participants (see Appendix - Supplementary Material). Herein, participants acknowledged reaching maximal speed and sustaining the highest speed possible throughout the 3MT without looking at the watch.

### Sessions 2 & 3

For sessions 2 and 3, at least 48 hours after session 1, ran at similar times of the day, and separated by at least 48 hours (maximal one week), participants performed a time-to-exhaustion test (TTE) on a level (TTE_Level_) and an inclined treadmill (TTE_Incline_). Participants refrained from heavy exercise one day prior to the testing and maintained their normal healthy dietary routine. Both sessions entailed the exact same procedures except for the treadmill’s incline, and we randomly assigned the order of these sessions in counterbalance, which the participants performed within one week. We used Stryd’s Indoor Treadmill Mode for both TTEs.

Following height and mass measurements and foot pod installation, we placed a near-infrared spectroscopy muscle oxygenation sensor (Moxy Monitor, Fortiori Design LLC, Hutchinson, Minnesota, USA) on the participant’s left and right rectus femoris muscles, halfway between the anterior superior iliac spine and the top part of the patella (Feldmann et al. 2019). We used the VO2 Master Manager App (Version 0.41.0, VO2 Master Health Sensors Inc., Vernon, BC, Canada), which allows the sampling of up to three Moxy Monitors for data recording. After a self-paced 8-minute warmup, participants performed two initialization trials of maximal 1 minute each, first on a level treadmill (0°) and after a 5-minute passive rest on the same treadmill inclined to 2.86° (5% grade) (Treadmetrix, Park City, UT, USA). The initialization trials determined the treadmill speeds to match the power and prepared the participants for the TTEs. The first initialization trial was set to the participant’s CS. The second initialization trial corresponded to the speed at which the participants averaged the same running power as within the first initialization trial. Following another passive rest of 5 minutes, participants ran at either CS for the TTE_Level_ on a 0° level treadmill or up a 2.86° incline on the same treadmill at the speed derived from the second initialization trial, matching the same power output of the first initialization trial for the TTE_Incline_ until exhaustion. We used CS as the reference marker instead of CP because Stryd’s Indoor Mode does not account for their reported Air Power, which estimates the air resistance at a given speed, considering the wind conditions affecting Stryd’s running power estimate outdoors. According to Stryd’s white paper, this can lead to a power difference of up to 10 W or more between Outdoor and Indoor Mode (Stryd Team 2020). Moreover, during our pilot testing phase, participants ran at 105% CP for both TTEs, for which the speed had to be increased to around 110% CS from the 3MT.

The following criteria standardized the end time of each TTE: a) Participants asked or gestured the researchers to stop the treadmill; b) Participants grabbed the safety railing of the treadmill unable to keep up the speed; c) Participants drifted backward beyond a zone we had marked on the treadmill with tape for more than 3 seconds; d) Researches stopped the treadmill as the participants showed severe signs of fatigue (e.g. losing balance, slipping) for safety, or e) Researches ended the test after 30 minutes of elapsed time. During the subsequent testing, for 26 out of the 28 total trials the participants asked the researchers to stop the treadmill (criterion a). We stopped the treadmill for one trial because the participant was drifting out of the marked zone for more than 3 seconds (criterion c); for another trial, we stopped the treadmill for safety as the participant showed severe signs of fatigue (criterion d). We measured RPE in 3-minute intervals (Scherr et al. 2013) (Borg_3min_, Borg_6min_, and so on) utilizing the 6-20 original Borg-Scale (Borg and Dahlström 1962). Additionally, examiners asked for RPE ratings at random intervals to keep the participants unaware of the elapsed time. These additional ratings were not considered for the analysis.

### Data Analysis

We used MATLAB (Version R2023b, The MathWorks Inc., Natick, MA, USA) for data and statistical analysis. The primary outcome variable was the time of both TTEs (Time_TTE_). Because of initial treadmill acceleration and power fluctuations were pronounced towards the end of each TTE, we defined Time_TTE_ as the total time [s] spent above a power threshold [W] two standard deviations lower than the participant’s mean power [W] from 30 seconds into the TTE until 10 seconds before the first NaN (Not a number) power reading. This enabled standardization without the subjectivity of hand-clocked stopwatch times between participants and examiners. We also calculated the average trial power (Power_TTE_) for each TTE using the same interval as for Time_TTE_.

We collected and exported %SmO_2_ from both Moxy sensors at 1 Hz (Matthews et al. 2023) and averaged sensor data of the two monitors. We removed a %SmO_2_ data set from the analysis if it contained more than 5% of NaN data or entailed more than two jumps of 5% between two consecutive data points over the entire trial length. We then averaged the subsequent %SmO_2_ raw data set over 10 seconds (Kirby et al. 2021; Matthews et al. 2023). We calculated the average trial %SmO_2_ as the mean %SmO_2_ over the entire exercise trial and the end trial %SmO_2_ as the final 5% of the time of each TTE (Kirby et al. 2021). We calculated the “delayed” slope (Δ%SmO_2_·Δmin^-1^) from data obtained 60 seconds after the start (beyond the onset phase) to the end of each trial (Kirby et al. 2021).

We estimated the metabolic rate [W·kg^-1^] during both TTEs using the RE3 equation for the TTE_Level_ using average trial speed [ᵄF = ᵃ6ᵄF] for level running (eq. 1; Looney et al. 2023) and the HTK equation for uphill running for the TTE_Incline_ (eq. 2; Hoogkamer et al. 2014; HTK based on the last name initials of the authors):

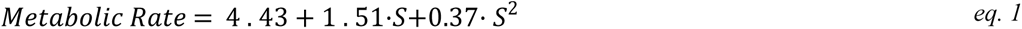

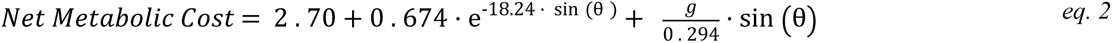

For calculating the gross metabolic rate of the TTE_Incline_, we multiplied by speed and added the standing metabolic rate of 1.53 W·kg^-1^ as measured in Hoogkamer et al. (2014).

### Statistical Analysis

We performed paired t-tests for Time_TTE_, Power_TTE_, Average Trial %SmO_2_, End Trial %SmO_2_, delayed slope, RPE (Borg_3min_, Borg_6min_, Borg_9min_, and Borg_End_), and estimated metabolic rate between the TTE_Level_ and the TTE_Incline_. Additionally, we compared CS and CP from the Stryd foot pod with CS_Coros_ and CP_Coros_ using paired t-tests. We included mean RPE time points of the TTE_Level_ and the TTE_Incline_ if more than ten paired observations were recorded. We excluded %SmO_2_ data for two participants from the statistical analysis according to the criteria listed above. An alpha level of 5% was set a priori.

## Results

### 3MT CS and CP

CS and CP during the 3MT were 4.65 ± 0.41 m·s^-1^ and 353 ± 83 W, respectively, for the Stryd foot pod. CS_Coros_ and CP_Coros_ were 4.99 ± 0.52 m·s^-1^ and 371 ± 93 W for Coros, both significantly higher than the results from the Stryd foot pod (p < 0.001 and p < 0.01, respectively). Fig. 1 details the Stryd’s and Coros’ speed and power curves of the 3MT for all participants.

**Fig. 1.**
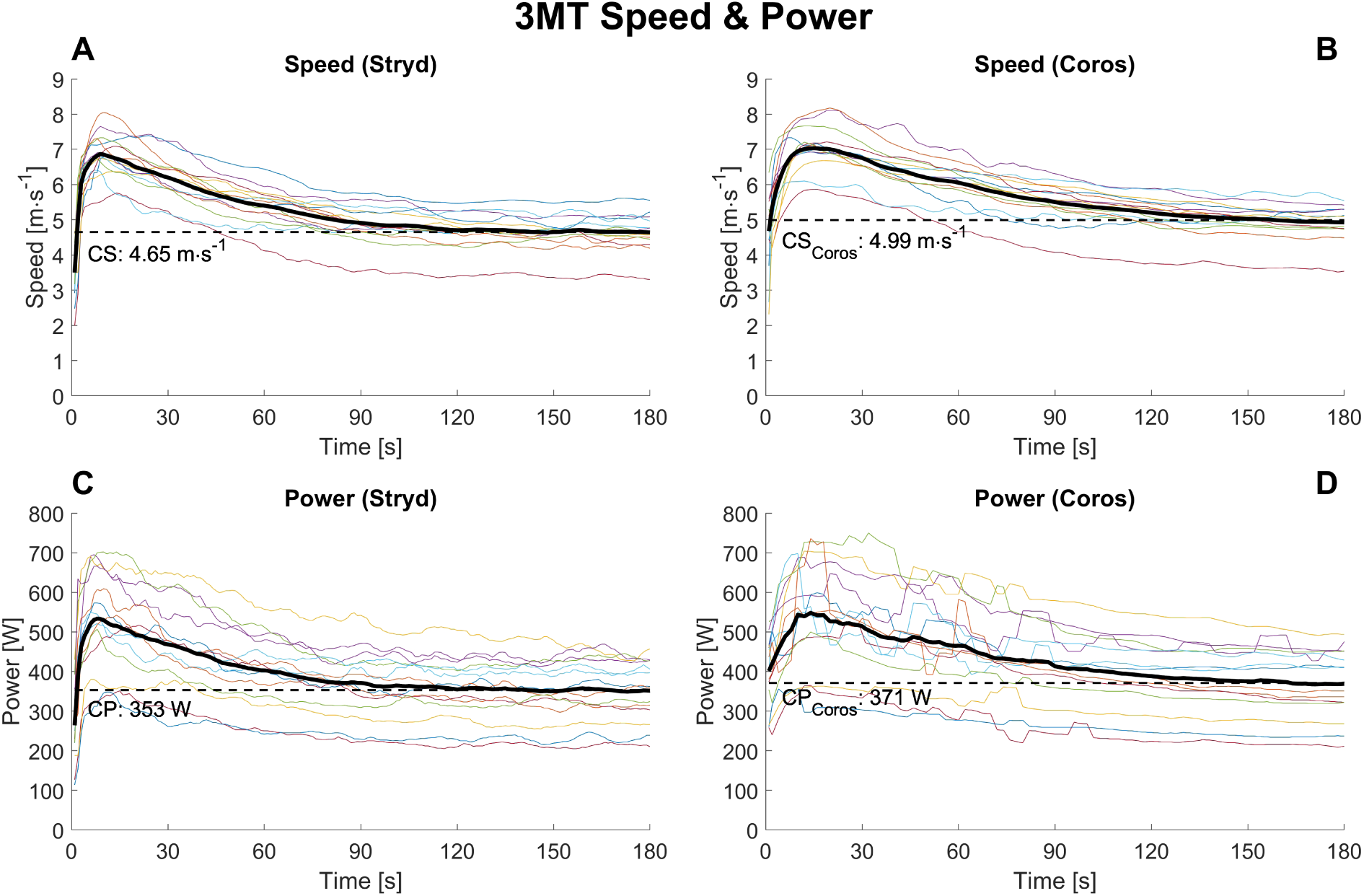
Speed [m·s^-1^] and power [W] recording of the Stryd foot pod (Fig. 1a and 1c) and the Coros Apex 2 watch (Fig. 1b and 1d). The black line represents the averaged speed and power data across participants. The dashed line represents the participants’ averaged CS, CP, CS_Coros_, and CP_Coros_. Each color represents one participant’s data

### Time_TTE_, Power_TTE_, and Speed

All 14 participants performed both TTEs. The participants ran 397 s (-42%) shorter in TTE_Level_ (539 ± 160 s) than in TTE_Incline_ (936 ± 336 s) (p < 0.001; Fig. 2). Power_TTE_ was similar between the trials with 331 ± 73 and 329 ± 68 W, respectively (p = 0.26). The average speed was 4.65 ± 0.41 m·s^-1^ for the TTE_Level_ and 3.51 ± 0.35 m·s^-1^ for the TTE_Incline_. Participants ran at 94.1 ± 3.3 and 93.8 ± 3.6 % of their CP during the TTE_Level_ and the TTE_Incline_ (p = 0.56).

**Fig. 2.**
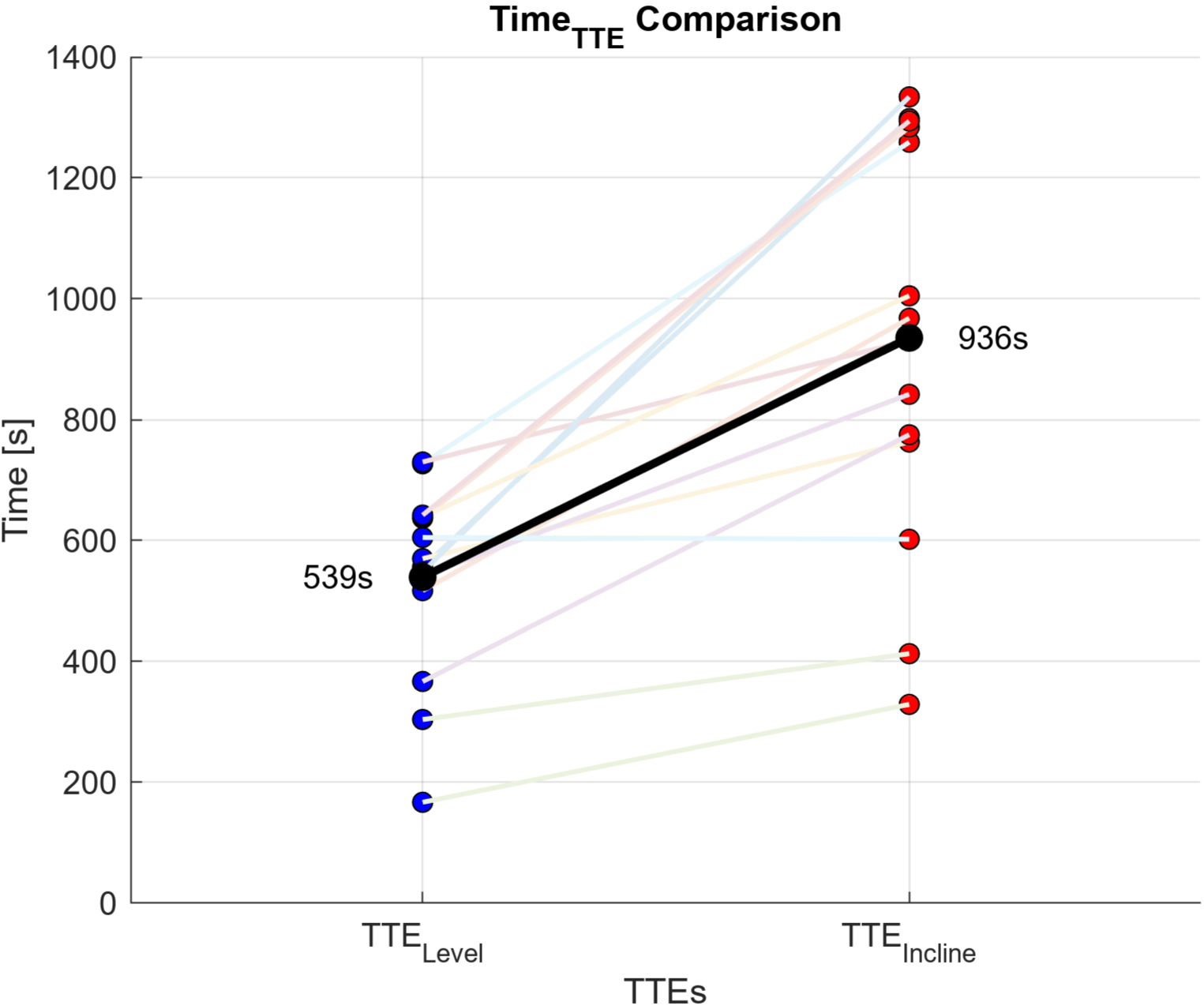
The participants ran substantially shorter in the level time-to-exhaustion test (TTE_Level_) than in the uphill time-to-exhaustion test (TTE_Incline_). The black line represents the averaged group data

### Muscle O_2_ saturation

Average trial %SmO_2_ for the TTE_Level_ was 33.3 ± 10.9 % compared to 37.9 ± 11.5 % for the TTE_Incline_ (p < 0.01), with an end trial %SmO_2_ of 21.3 ± 10.7 % vs. 29.3 ± 11.4 % (p < 0.001), respectively. The delayed slope for the TTE_Level_ was -2.48 ± 1.32 %·min^-1^ compared to -1.62 ± 2.11 %·min^-1^ for the TTE_Incline_ (p < 0.05). Fig. 3 displays the SmO_2_ of the TTE_Incline_ and the TTE_Level_ normalized to 100 data points.

**Fig. 3.**
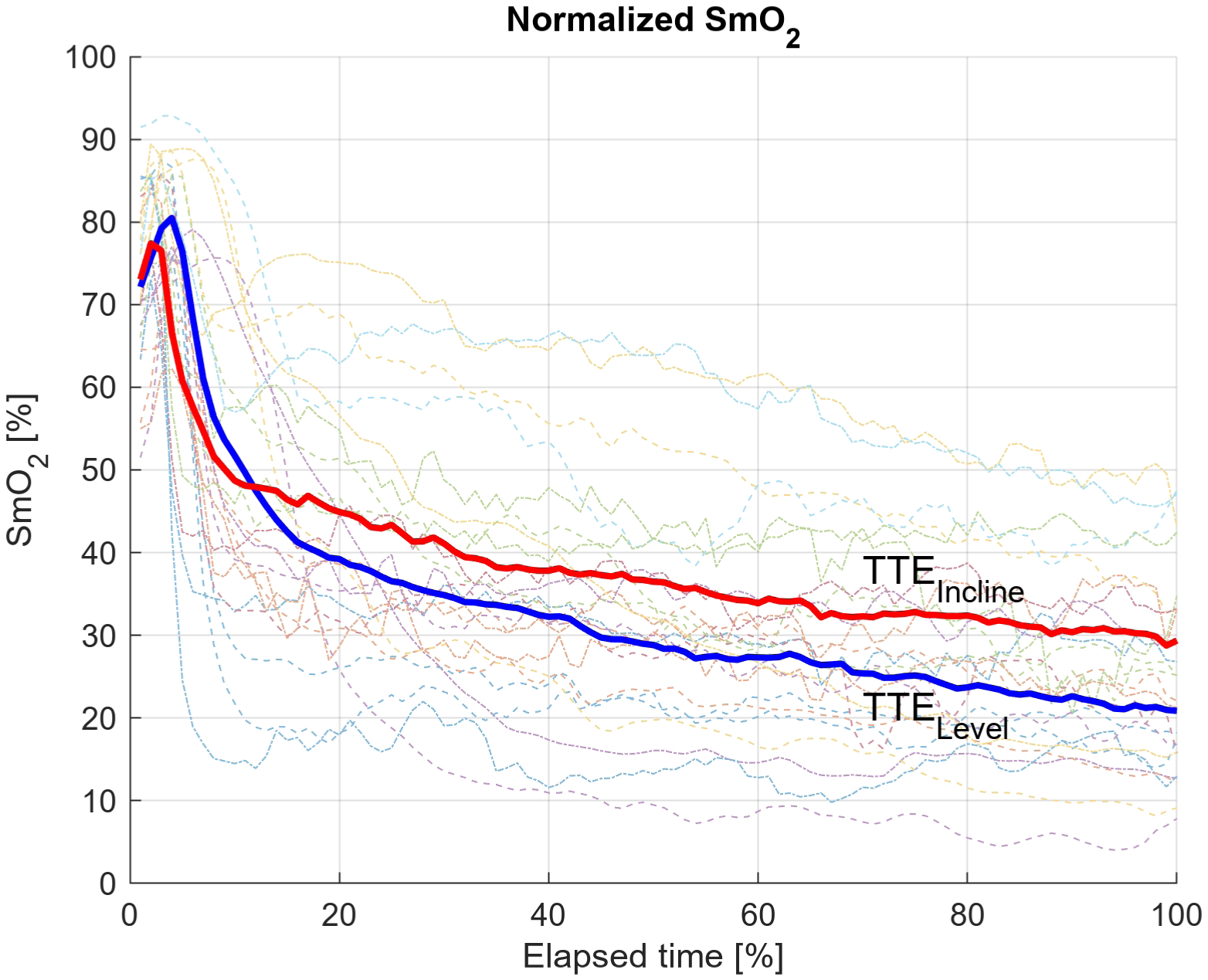
Muscle oxygenation values (SmO_2_ [%]) normalized to trial time decreased relatively faster and more during the level time-to-exhaustion test (TTE_Level_) than during the uphill time-to-exhaustion test (TTE_Incline_). The bold blue line represents the group’s averaged %SmO2 data for the TTE_Level_ and the bold red line the group’s averaged %SmO2 data for the TTE_Incline_. Individual data for the TTE_Level_ and TTE_Incline_ are matched in color (TTE_Level_: Dashed line; TTE_Incline_: dash-dotted line)

### RPE (Borg)

Tab. 1 contains a summary of the RPE values at the analyzed time points. Borg_End_ for the TTE_Level_ was 19.5 ± 0.7, similar to 19.4 ± 0.8 for the TTE_Incline_ (p = 0.34). For both trials, all participants reported a Borg_End_ of 18 or more. At each of the 3-minute intervals, the Borg values were significantly higher for TTE_Level_ than for the TTE_Incline,_ except for Borg_End_.

**Tab. 1.**
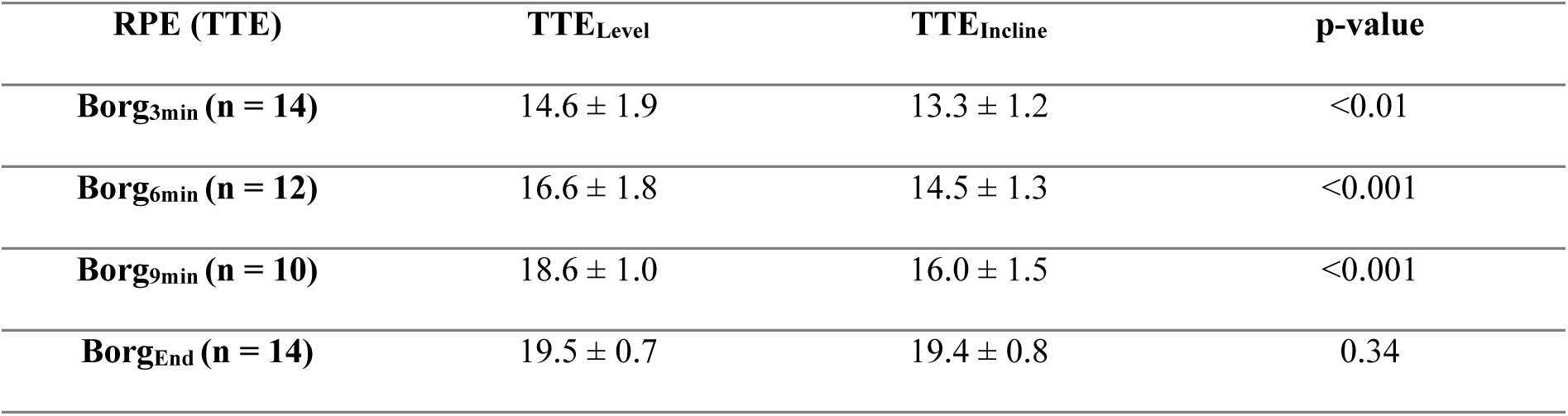
Rating of perceived exertion of the time-to-exhaustion tests (TTEs) using the 6-20 Borg Scale assessed at 3, 6, and 9-minute elapsed time was higher during the level TTE than during the uphill TTE at matched time points, but similar at the end of each TTE (Borg_End_)

### Estimated Metabolic Rate

The estimated metabolic rate based on running speed and slope was significantly higher during TTE_Level_ (19.51 ± 1.99 W·kg^-1^) than during TTE_Incline_ (17.82 ± 1.62 W·kg^-1^) (p < 0.001; Fig. 4).

**Fig. 4.**
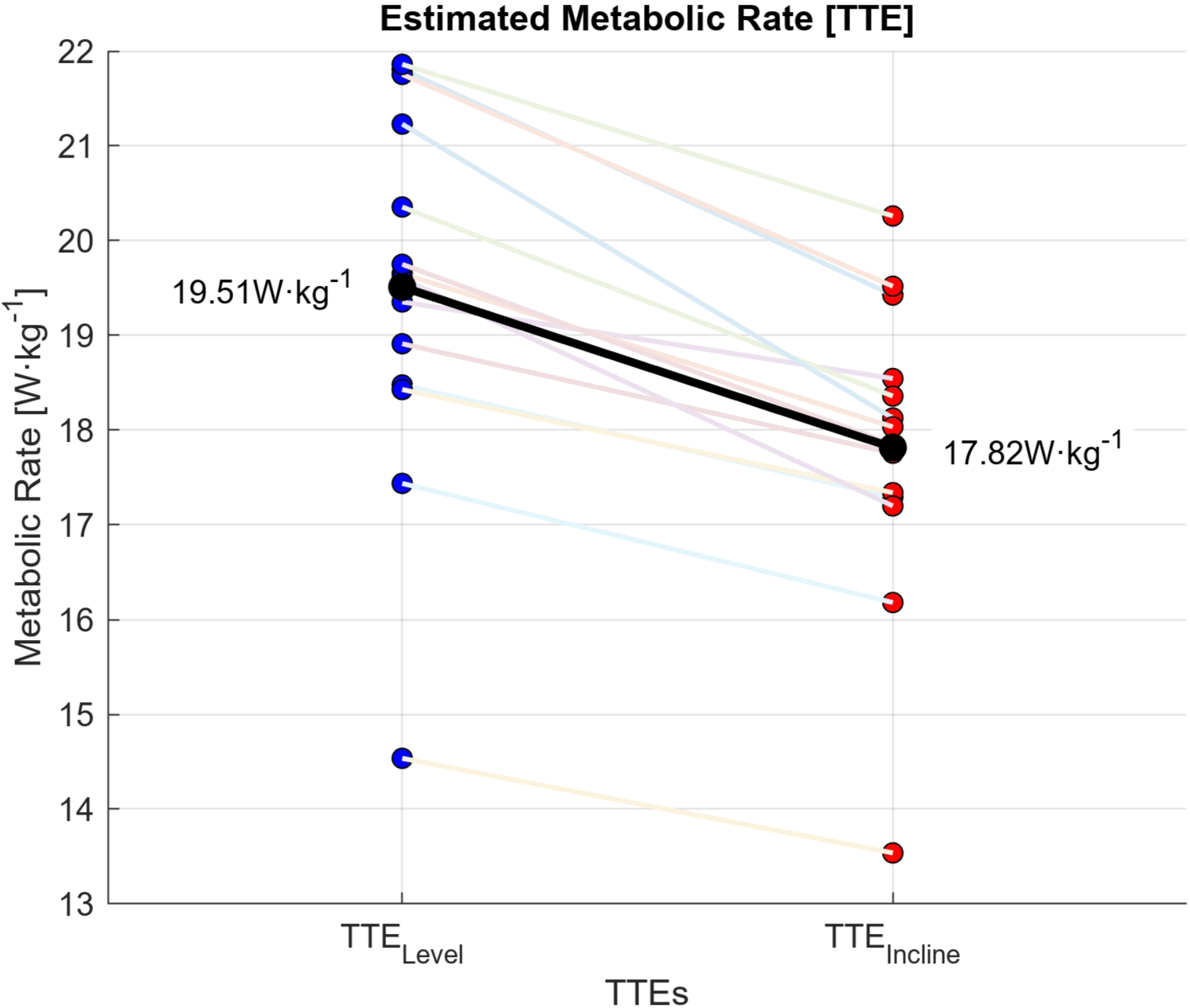
Using the average speed of both time-to-exhaustion trials, the estimated metabolic rate [W·kg^-1^] was higher for the level time-to-exhaustion test (TTE_Level_) than for the uphill time-to-exhaustion test (TTE_Incline_). Individual metabolic rates are connected by a colored solid line. The black line represents averaged group data

## Discussion

Running to exhaustion at matching power outputs using Stryd’s foot pod resulted in significantly shorter Time_TTE_ during level running than uphill running. This was accompanied by higher muscle oxygen demands measured by the near-infrared spectroscopy sensors, perceived effort (RPE), and estimated metabolic rate during the TTE_Level_ than during the TTE_Incline_.

Our findings imply that when exercising at a comparable effort between the level and uphill trials as assessed by Stryd running power, one actually performs at a lower intensity during uphill running. From a practical perspective, estimating running power mainly adds value beyond running speed when used for altered conditions, such as uphill running or running into a headwind. Stryd running power has been validated against metabolic rate across a wide range of speeds in level running (Imbach et al. 2020; Taboga et al. 2022). We did not directly measure metabolic rate, but using existing empirical equations for metabolic rate during level and uphill running, we estimated significantly higher metabolic rates at CS for the level running trial. We focused on CS and CP as the boundaries of the heavy to the severe exercise intensity domain (Poole et al. 2016) to ensure comparability between both trials at an intensity representative for race performances and critical intensity-based workouts (Clark et al. 2013; Figueiredo et al. 2023).

To adjust the gradient for the TTE_Incline_ in the Stryd app, we used the app’s Indoor Mode for both TTEs. For outdoor running, Stryd assesses elevation from barometric pressure changes to correct their power estimation (Stryd Outdoor Mode), but as barometric pressure changes do not occur during uphill treadmill running, a runner needs to manually set the treadmill grade in the app in Indoor Mode. We conclude that the incline factor of Stryd’s Indoor Mode does not sufficiently capture the alteration in mechanical power of uphill running.

We estimated the metabolic rate of the TTE_Level_ and the TTE_Incline_ and found significantly higher values for the TTE_Level_ (19.51 ± 1.99 W·kg^-1^) compared to the TTE_Incline_ (17.82 ± 1.62 W·kg^-1^) at matching running power outputs. We used the HTK equation (Hoogkamer et al. 2014) to estimate the metabolic rate for the TTE_Incline_ and the RE3 equation (Looney et al. 2023) for the TTE_Level_. It is important to note that the HTK equation was developed with speeds of up to 3 m·s^-1^, yet our study’s average TTE_Incline_ speed was 3.51 m·s^-1,^ with only one participant running slower than 3 m·s^-1^ for this trial. Recently, the authors updated their equations (Looney et al. 2025), so we recalculated the metabolic estimates using these new equations as well. For both updated equations, metabolic rate was higher during the TTE_Level_ than for the TTE_Incline_. With the updated HTK equation, the estimated metabolic rate was 18.88 ± 1.56 W·kg^-1^ during the TTE_Level_ vs. 18.44 ± 1.70 W·kg^-1^ during the TTE_Incline_ (p < 0.01), and with the updated RE3 (now with an uphill term), the estimated metabolic rate was 19.51 ± 1.99 W·kg^-1^ during the TTE_Level_ vs. 18.59 ± 1.82 W·kg^-1^ during the TTE_Incline_ (p < 0.001). We did not measure metabolic rate because we could not assume aerobic steady state at CS (Poole et al. 1988), also indicated by the negative muscle oxygenation slope we observed during the TTEs. We were particularly interested in assessing the usability of the Stryd foot pod to assess running power at critical intensity at various slopes; future studies could compare metabolic rate between level and uphill running at matched running power at lower, aerobic steady-state intensities.

Based on these differences in metabolic rate estimations between level and uphill running at similar Stryd running power, we then decided to estimate the metabolic efficiency for each trial by dividing the estimated metabolic power by the recorded running power. For level running, the estimated metabolic efficiency was around 24%, independent of body mass (Fig. 5a). For uphill running, the estimated metabolic efficiency decreased as a function of body mass from 29% for the least heavy participant (43.1 kg) down to 24% for the heaviest participants (over 90 kg) (Fig. 5a). To further assess the metabolic efficiency during level running, relative to the estimated metabolic rate, we created a Bland-Altman plot (Fig. 5b), which confirmed the consistency of this efficiency estimate for level running.

**Fig. 5.**
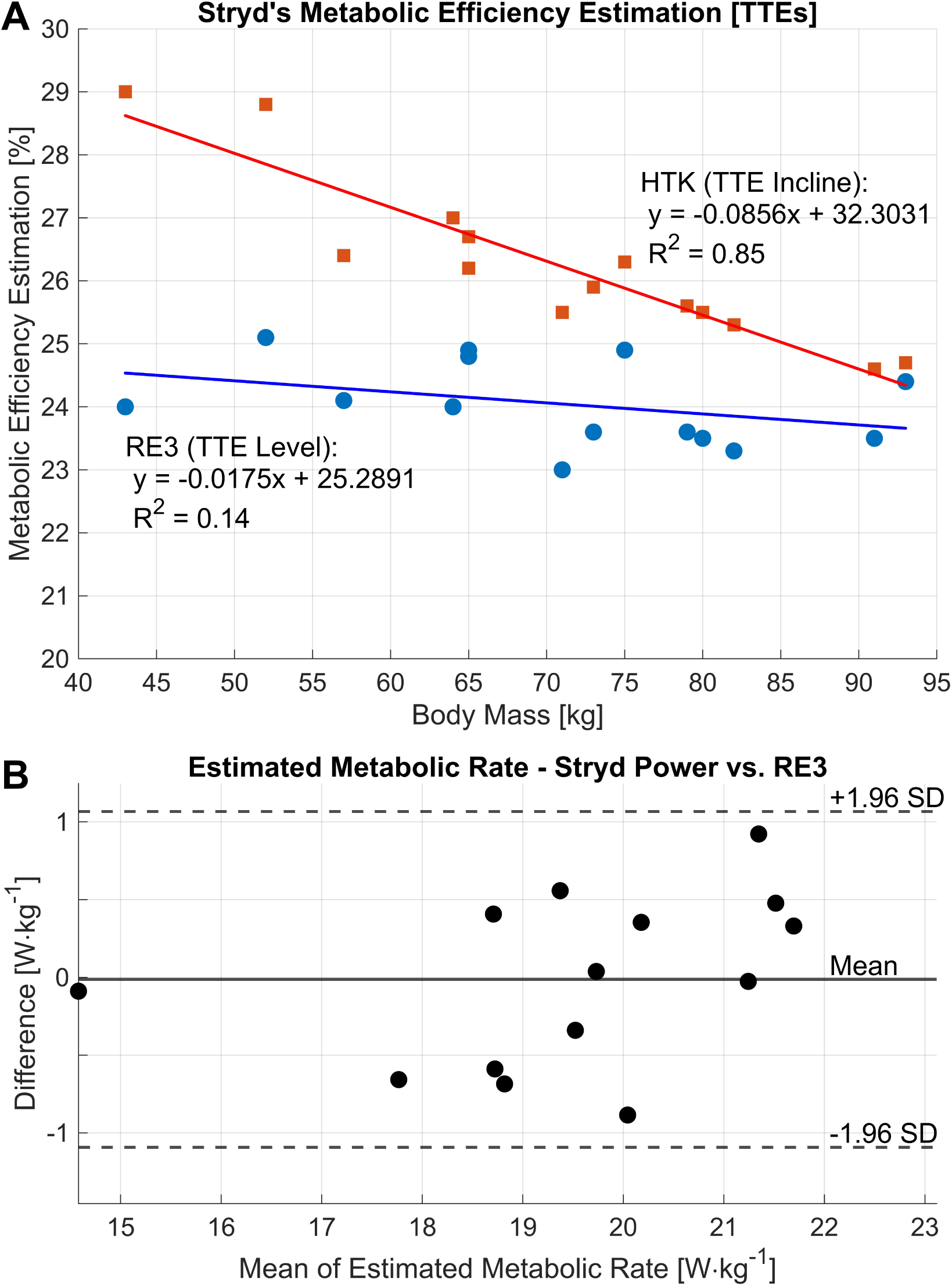
Metabolic efficiency estimation using linear regression for the level time-to-exhaustion (TTE_Level_ in blue) showed a weak but high correlation for the uphill time-to-exhaustion test (TTE_Incline_ in red) for body mass (Fig. 5a). Linear regression equations and R^2^ are displayed next to the regression curves. For the TTE_Level_, we used the RE3 equation (Looney et al. 2023), and for the TTE_Incline_, we used the HTK equation (Hoogkamer et al. 2014) and corresponding running power of the time-to-exhaustion test (Power_TTE_). The estimated metabolic rate of the level time-to-exhaustion test (TTE_Level_) was similar between Stryd-derived Power vs. Speed from the TTE_Level_ using the RE3 equation and a 24% metabolic efficiency (Fig. 5b)

We could not measure Coros Power during the TTEs as the Coros Watch does not estimate power for treadmill running. Therefore, we cannot compare the running power of the TTEs between devices. During the 3MT, Coros Power visually fluctuated more than Stryd Power (Fig. 1). This might be due to Coros’ power estimation derived from a more variable GPS-based speed. Also, the recorded total distance during the 3MT using the Coros watch in track mode in lane 1 did not reliably match the actual covered distance on the track, typically overestimating the total distance.

As detailed in the method section, we used CS for the TTE_Level_ and matching power output for the TTE_Incline_ instead of CP because of differences between overground (track) and treadmill running, which affect the runner’s speed-power ratio. During the TTE_Level_, we observed a minor power drift of 2.19 ± 1.82 % from the start of the TTE (average over 0:30-1:00 min) to the end of the TTE (average over 0:40 to 0:10 min before the first NaN reading). For the TTE_Incline_, the drift was not significant (0.17 ± 0.90 %). We chose these time windows to capture the power output while running at CS without the initial acceleration and deceleration phase of the TTE. Even though this power drift is small compared to accuracy numbers reported for a range of cycling power meters (1-5%) (Maier et al. 2017), we wanted to see if it was related to changes in cadence during the TTEs. We detected cadence drift of -2.99 ± 1.64% for the TTE_Level_ and -1.96 + 2.29% for the TTE_Incline_ (p < 0.05), yet the power drift was not correlated to the cadence drift using linear regression analysis (R^2^ = 0.02). Participants averaged 182.0 ± 10.1 steps per minute for the TTE_Level_ and 173.1 ± 9.5 steps per minute for the TTE_Incline_.

We assessed muscle oxygen demand during the TTEs and observed significantly higher oxygen depletion for the TTE_Level_ over time using the delayed slope concept (Kirby et al. 2021). The oxygen level decreased by 2.48 ± 1.32 %·min^-1^ for the TTE_Level_ and by 1.62 ± 2.11 %·min^-1^ for the TTE_Incline_. Kirby et al. (2021) reported a delayed slope of -2.65 ± 0.77 %·min^-1^ during exercise within the severe intensity domain and a consistently positive slope for exercise in the heavy intensity domain of 0.53 ± 0.14 %·min^-1^. These findings suggest that our participants likely performed within the severe intensity domain, especially for the TTE_Level_. This might explain the relatively short Time_TTE_ we observed for both trials running at critical intensity, the intensity which theoretically can be sustained indefinitely, but in reality for approximately 16 to 30 minutes (Pepper et al. 1992; Bull et al. 2008; Figueiredo et al. 2021). The non-competitive nature of the lab setting and the unawareness of participants of elapsed time and running speed might have contributed to the relatively short Time_TTE_ we observed, although our Time_TTE_ was still shorter than reported by Pepper et al. (1992) in similar lab and awareness conditions (986 ± 365 seconds) who determined CS from a series of 5 TTEs on a level treadmill instead of using a 3MT.

Numerous papers have compared the 3MT-derived CS or CP estimations to the current gold standard of time trial efforts in running (e.g., Broxterman et al. 2013; Gama et al. 2018; Hunter et al. 2023) and have demonstrated similar CS and CP estimations on a group level. However, critical intensity estimations can substantially differ on an individual level, as shown by Broxterman et al. (2013). On a group level, the difference between methods averaged out at only 2.32% for Broxterman’s cohort, but importantly, the average absolute intra-individual difference was 6.58%, ranging from 0.52% up to 19.20% among participants. Such an over- and underestimation of the 3MT-derived critical intensity will substantially impact a runner’s time-to-exhaustion because of the hyperbolic relationship between running speed to time-to-exhaustion (Daniels et al. 1978; Poole et al. 2016). For example, the current men’s world records in 5000 m (12:35.36 min) and 10.000m (26:11.00 min) set by Joshua Cheptegei in 2020 differ only 3.8% in running speed, but this small speed difference enables an effort more than twice as long in duration. For our participants, the estimated metabolic rate was ∼9% lower during the TTE_Incline_, and they managed to run 42% longer on average, in line with this hyperbolic relationship.

Importantly the TTE durations varied substantially between individuals (167 s to 730 s), which could have been related to over- or underestimations of their 3MT-derived critical intensity. Therefore, we evaluated the execution of 3MT using existing criteria (Muniz-Pumares et al. 2019). The 3MT assumes that towards the end of the test, a runner can only sustain their CS or CP as their finite capacity above CS or CP - called D-Prime (D’) or W-Prime (W’) (proxies for anaerobic working capacity) - has been fully depleted (Pettitt 2016) while the mean speed or power of the last 30 seconds often serves as CS or CP approximation (Pettitt 2016). Muniz-Pumares et al. (2019) developed criteria for the 3MT in cycling to assess whether a participant has properly executed their test. Hunter et al. (2023) used these criteria to establish running CS and CP values based on Stryd’s foot pod. These criteria include: 1) a plateau in power output in the last 30 seconds of the test; 2) the attainment of peak power output within the first 10 seconds of the test; 3) rapid initial decrease of power output to deplenish >90% of the work above end power, a proxy for anaerobic working capacity; 4) no decrease in power output of >5% of end power for >5s during the test; 5) an end test cycling cadence within 10 rpm of preferred cadence; 6) the attainment of VO_2_max, and 7) a blood lactate concentration >8mmol·L^-1^. We have assessed 4 out of the 7 criteria (criteria 1 - 4) for the 3MT using the running power of the Stryd foot pod and observed 8.71 ± 4.34 s (Peak Power: 546 ± 122 W; Peak Speed: 6.95 ± 0.58 m·s^-1^) for criterion 2, 92.37 ± 4.54% for criterion 3, and 3.50 ± 5.26 s for criterion 4. Additionally, as there is currently no standardized criterion for evaluating the plateau during end power (criterion 1, Muniz-Pumares et al. 2019), we developed a standard to assess steady state during end power that assesses if any of the data points exceed 2 standard deviations above or below the mean. For our 3MT, 4 out of 14 participants met all criteria, while criterion 1) was met by 11 out of 14 participants, criterion 2) by 11 out of 14 participants, criterion 3) by 10 out of 14 participants, and criterion 4) by 11 out of 14 participants. Herein, 12 out of 14 participants missed only one criterion, 1 out of 14 missed two criteria, and 1 out of 14 missed three criteria. Interestingly, the participant who missed three criteria ran the second longest TTE_Level_ of all the participants with 727 seconds. We cannot analyze criterion 5 as it is specific to cycling and cannot be applied directly to running, and criteria 6 and 7 require additional equipment. Compared to Hunter et al. (2023) who assessed two 3MTs, our participant cohort reached a higher peak speed (6.95 ± 0.58 m·s^-1^ vs. 5.92 ± 1.09 and 5.69 ± 1.04), and a higher peak power (546 ± 122 W vs. 451 ± 101 W and 454 ± 106 W), which they also reached after a shorter time (8.71 ± 4.34 s vs. 12.6 ± 3.6 s and 17.0 ± 10.8 s) (criterion 2). Moreover, a higher percentage of our participants met criterion 3 (71.4% vs. 21.7% and 30.4%) and depleted their W’ by more than >90% within the first 90 seconds. However, 3 out of our 14 participants missed criterion 4 and dipped below 5% of their end power for more than 5 seconds, compared to none of the participants in Hunter’s group. Still, these participants did not run shorter than the other participants in the TTE_Level_. These findings suggest that our participants may have depleted their anaerobic working capacity at a faster rate than Hunter’s group, as demonstrated by criterion 3, but dipped below their end power and might have, therefore, recuperated some of their finite working capacity that would elevate their end power. We conclude that standardized criteria for the 3MT, as developed by Muniz-Pumares et al. (2019) for cycling, would enhance the comparability of the 3MT for running across studies. However, we need to acknowledge the differences between stationary bike cycling and overground running, particularly in the acceleration phase.

### Limitations (261w)

We used Stryd’s foot pod on the track using the Stryd app’s Outdoor Mode and on the treadmill using the Stryd app’s Indoor Mode. Our results are mainly based on using Stryd’s Indoor Mode on the treadmill. We must acknowledge the differences between Stryd’s Indoor Mode and Stryd’s Outdoor Mode in the way they consider elevation changes to estimate running power. Therefore, conclusions on the usability of Stryd’s food pod for running at critical intensity can only be made for treadmill uphill running. Further, we performed the 3MT in track lane 1, and as such, participants were aware of how much distance they had covered, which could have affected the 3MT results. We decided to use lane 1 to enhance the Coros GPS watch accuracy since, during pilot testing, we identified substantial differences in the participant’s running distance with the Coros Watch captured distance when using lane 5 or 6 (input in the Coros’ track mode setting). In hindsight, this anomaly seems to be independent of the running lane. As discussed earlier, we did not measure metabolic rate but estimated it using literature-based equations (HTK; Hoogkamer et al. 2014 and RE3; Looney et al. 2023) because we could not assume aerobic steady state at CS, and those estimations are likely not to be entirely accurate at the individual level. Lastly, we did not fully control for diet and training leading into the TTEs, but randomized the level and uphill trial order in counterbalance, and it is unlikely that our strong findings were solely due to varying diet or training status.

### Perspective

We provide the first scientific work that examined running power on distinct incline levels at threshold intensity. Previous research has shown a strong linear correlation between running power and metabolic rate for level running over a range of different running velocities (Imbach et al. 2020, Taboga et al. 2022). However, running power lacks validity to capture changes in metabolic demand during altered conditions other than running velocity, such as when using advanced footwear technology vs. traditional running shoes (Hudgins et al. 2024), and non-level running conditions have been overlooked in the current literature (Apte et al., 2023). Our findings suggest that uphill running power, as observed during treadmill running using the Stryd foot pod, does not serve as a viable metric to quantify a runner’s metabolic effort. Moreover, we demonstrate that running power estimations between foot- vs. wrist-based devices can significantly differ. Therefore, runners should use running power with caution to pace and guide their training or racing, especially in non-level running conditions.

## Conclusion

Our data suggests that athletes should be careful when using running power at critical intensity to guide their racing and training in non-level conditions. Further, our findings suggest that running power significantly differs between foot- vs. wrist-based wearables.

## Supporting information

Instructional Script for the 3-minute all-out test

Original Data of the testing

## Author Contributions

Nikolai Kirner (N.K.) and Wouter Hoogkamer (W.H.) conceived and designed research; N.K. performed experiments; N.K. and W.H. analyzed data; N.K. and W.H. interpreted results of experiments; N.K. prepared figures; N.K. and W.H. drafted the manuscript; N.K. and W.H. edited and revised the manuscript; N.K. and W.H. approved the final version of the manuscript.

## Acknowledgments

We thank all runners for participating in the study and Caroline Kolmodin, Keegan Butler, Nadia Surin, and Nathan Hammerschmitt Le Gal for their help with data collection. We also thank Dr. Fabian Stöcker for providing the Stryd foot pod.

## Declarations

### Ethical Approval

Approval was obtained from the Institutional Review Board of the University of Massachusetts Amherst (Protocol Nr.: 5639). The procedures used in this study adhere to the tenets of the Declaration of Helsinki.

### Informed Consent

Informed consent was obtained from all individual participants included in the study.

### Data availability

The datasets generated during and/or analyzed during the current study are available from the corresponding author on reasonable request.

## Abbreviations

3MT: 3-minute all-out test
Borg_3min_: Rating of Perceived Exertion using the Borg scale at 3-min elapsed time
Borg_6min_: Rating of Perceived Exertion using the Borg scale at 6-min elapsed time
Borg_9min_: Rating of Perceived Exertion using the Borg scale at 9-min elapsed time
Borg_End_: Rating of Perceived Exertion using the Borg scale at the end of the test
CP / CP_Coros_: Critical Power / Critical Power derived from the Coros Watch
CS / CS_Coros_: Critical Speed / Critical Speed derived from the Coros Watch
D’: D-Prime (a proxy for anaerobic working capacity above Critical Speed)
HTK: Hoogkamer et al.’s (2014) equation for estimating metabolic rate
NaN: Not a number
Power_TTE_: Running power of the time-to-exhaustion test
RE3: Looney et al.’s (2023) equation for estimating metabolic rate
RPE: Rating of Perceived Exertion
Time_TTE_: Time of the time-to-exhaustion test
TTE: Time-to-exhaustion test
TTE_Incline_: Uphill time-to-exhaustion test
TTE_Level_: Level time-to-exhaustion test
W’: W-Prime (a proxy for anaerobic working capacity above Critical Power)

