## Supplementary material for "Wearable Assessment of Level and Uphill Running at Critical Intensity": Instructional Script for the 3-minute all-out test

Standardized instructional script for the 3-minute all-out test (“Robber Analogy”):

*Imagine you have just gone to a bank to withdraw some cash. Suddenly, two robbers' storm in and take everyone hostage, including you. You notice the door is open and muster all your courage to make a run for it. One of the robbers sees you and starts chasing you with a knife as you dash towards the police station, which is just three minutes away.*

*You sprint as fast as you can, knowing that the first 200 meters are crucial to avoid being caught. After that, you do your best to maintain your speed, determined not to let the robber catch up. There is no time to check the clock, as any distraction could slow you down. Even though you start to tire before reaching the police station, you push yourself to keep up the pace, driven by the hope of saving the others.*
